# A COVID Moonshot: assessment of ligand binding to the SARS-CoV-2 main protease by saturation transfer difference NMR spectroscopy

**DOI:** 10.1101/2020.06.17.156679

**Authors:** Anastassia L. Kantsadi, Emma Cattermole, Minos-Timotheos Matsoukas, Georgios A. Spyroulias, Ioannis Vakonakis

## Abstract

Severe acute respiratory syndrome coronavirus 2 (SARS-CoV-2) is the etiological cause of the coronavirus disease 2019, for which no effective therapeutics are available. The SARS-CoV-2 main protease (M^pro^) is essential for viral replication and constitutes a promising therapeutic target. Many efforts aimed at deriving effective M^pro^ inhibitors are currently underway, including an international open-science discovery project, codenamed COVID Moonshot. As part of COVID Moonshot, we used saturation transfer difference nuclear magnetic resonance (STD-NMR) spectroscopy to assess the binding of putative M^pro^ ligands to the viral protease, including molecules identified by crystallographic fragment screening and novel compounds designed as M^pro^ inhibitors. In this manner, we aimed to complement enzymatic activity assays of M^pro^ performed by other groups with information on ligand affinity. We have made the M^pro^ STD-NMR data publicly available. Here, we provide detailed information on the NMR protocols used and challenges faced, thereby placing these data into context. Our goal is to assist the interpretation of M^pro^ STD-NMR data, thereby accelerating ongoing drug design efforts.

## Introduction

Infections by the severe acute respiratory syndrome coronavirus 2 (SARS-CoV-2) resulted in approximately 1.8 million deaths in 2020 (1) and led to the coronavirus 2019 (COVID-19) pandemic (2–4). SARS-CoV-2 is a zoonotic betacoronavirus highly similar to SARS-CoV and MERS-CoV, which caused outbreaks in 2002 and 2012, respectively (5–7). SARS-CoV-2 encodes its proteome in a single, positive-sense, linear RNA molecule of ^~^30 kb length, the majority of which (^~^21.5 kb) is translated into two polypeptides, pp1a and pp1ab, via ribosomal frame-shifting (8, 9). Key viral enzymes and factors, including most proteins of the reverse-transcriptase machinery, inhibitors of host translation and molecules signalling for host cell survival, are released from pp1a and pp1ab via post-translational cleavage by two viral cysteine proteases (10). These proteases, a papain-like enzyme cleaving pp1ab at three sites, and a 3C-like protease cleaving the polypeptide at 11 sites, are primary targets for the development of antiviral drugs.

The 3C-like protease of SARS-CoV-2, also known as the viral main protease (M^pro^), has been the target of intense study owing to its centrality in viral replication. M^pro^ studies have benefited from previous structural analyses of the SARC-CoV 3C-like protease and the earlier development of putative inhibitors (11–14). The active sites of these proteases are highly conserved, and peptidomimetic inhibitors active against M^pro^ are also potent against the SARS-CoV 3C-like protease (15, 16). However, to date no M^pro^-targeting inhibitors have been validated in clinical trials. In order to accelerate M^pro^ inhibitor development, an international, crowd-funded, open-science project was formed under the banner of COVID Moonshot (17), combining high-throughput crystallographic screening (18), computational chemistry, enzymatic activity assays and mass spectroscopy (19) among the many methodologies contributed by collaborating groups.

As part of COVID Moonshot, we utilised saturation transfer difference nuclear magnetic resonance (STD-NMR) spectroscopy (20–22) to investigate the M^pro^ binding of ligands initially identified by crystallographic screening, as well as molecules designed specifically as non-covalent inhibitors of this protease. Our goal was to provide orthogonal information on ligand binding to that which could be gained by enzymatic activity assays conducted in parallel by other groups. STD-NMR is a proven method for characterising the binding of small molecules to biological macromolecules, able to provide both quantitative affinity information and structural data on the proximity of ligand chemical groups to the protein. Here, we provide detailed documentation on the NMR protocols used to record these data and highlight the advantages, limitations and assumptions underpinning our approach. Our aim is to assist the comparison of M^pro^ STD-NMR data with other quantitative measurements, and facilitate the consideration of these data when designing future M^pro^ inhibitors.

## Materials and Methods

### Protein production and purification

We created a SARS-CoV-2 M^pro^ genetic construct in pFLOAT vector (23), encoding for the viral protease and an N-terminal His_6_-tag separated by a modified human rhinovirus (HRV) 3C protease recognition site, designed to reconstitute a native M^pro^ N-terminus upon HRV 3C cleavage. The M^pro^ construct was transformed into *Escherichia coli* strain Rosetta(DE3) (Novagen) and transformed clones were pre-cultured at 37 °C for 5 h in lysogeny broth supplemented with appropriate antibiotics. Starter cultures were used to inoculate 1 L of Terrific Broth Autoinduction Media (Formedium) supplemented with 10% v/v glycerol and appropriate antibiotics. Cell cultures were grown at 37 °C for 5 h and then cooled to 18 °C for 12 h. Bacterial cells were harvested by centrifugation at 5,000 x g for 15 min.

Cell pellets were resuspended in 50 mM trisaminomethane (Tris)-Cl pH 8, 300 mM NaCl, 10 mM imidazole buffer, incubated with 0.05 mg/ml benzonase nuclease (Sigma Aldrich) and lysed by sonication on ice. Lysates were clarified by centrifugation at 50,000 x g at 4 °C for 1 h. Lysate supernatants were loaded onto a HiTrap Talon metal affinity column (GE Healthcare) pre-equilibrated with lysis buffer. Column wash was performed with 50 mM Tris-Cl pH 8, 300 mM NaCl and 25 mM imidazole, followed by protein elution using the same buffer and an imidazole gradient from 25 to 500 mM concentration. The His_6_-tag was cleaved using home-made HRV 3C protease. The HRV 3C protease, His_6_-tag and further impurities were removed by a reverse HiTrap Talon column. Flow-through fractions were concentrated and applied to a Superdex75 26/600 size exclusion column (GE Healthcare) equilibrated in NMR buffer (150 mM NaCl, 20 mM Na_2_HPO_4_ pH 7.4).

### Nuclear magnetic resonance (NMR) spectroscopy

All NMR experiments were performed using a 950 MHz solution-state instrument comprising an Oxford Instruments superconducting magnet, Bruker Avance III console and TCI probehead. A Bruker SampleJet sample changer was used for sample manipulation. Experiments were performed and data processed using TopSpin (Bruker). For direct STD-NMR measurements, samples comprised 10 μM M^pro^ and variable concentrations (20 μM – 4 mM) of ligand compounds formulated in NMR buffer supplemented with 10% v/v D_2_O and deuterated dimethyl sulfoxide (D_6_-DMSO, 99.96% D, Sigma Aldrich) to 5% v/v final D_6_-DMSO concentration. In competition experiments, samples comprised 2 μM M^pro^, 0.8 mM of ligand x0434 and variable concentrations (0 – 20 μM) of competing compound in NMR buffer supplemented with D_2_O and D_6_-DMSO as above. Sample volume was 140 μL and samples were loaded in 3 mm outer diameter SampleJet NMR tubes (Bruker) placed in 96-tube racks. NMR tubes were sealed with POM balls.

STD-NMR experiments were performed at 10 °C using a pulse sequence described previously (20) and an excitation sculpting water-suppression scheme (24). Protein signals were suppressed in STD-NMR by the application of a 30 msec spin-lock pulse. We collected time-domain data of 16,384 complex points and 41.6 μsec dwell time (12.02 kHz sweepwidth). Data were collected in an interleaved pattern, with on- and off-resonance irradiation data separated into 16 blocks of 16 transients each (256 total transients per irradiation frequency). Transient recycle delay was 4 sec and on- or off-resonance irradiation was performed using 0.1 mW of power for 3.5 sec at 0.5 ppm or 26 ppm, respectively, for a total experiment time of approximately 50 minutes. Reconstructed time-domain data from the difference of on- and off-resonance irradiation (STD spectra) or only the off-resonance irradiation (reference spectra) were processed by applying a 2 Hz exponential line broadening function and 2-fold zero-filling prior to Fourier transformation. Phasing parameters were derived for each sample from the reference spectra and copied to the STD spectra. ^1^H peak intensities were integrated in TopSpin using a local-baseline adjustment function. Data fitting to extract K_d_ values were performed in OriginPro (OriginLab). The folded state of M^pro^ in the presence of each ligand was verified by collecting ^1^H NMR spectra similar to Fig. 1A from all samples ahead of STD-NMR experiments.

**Figure 1:**
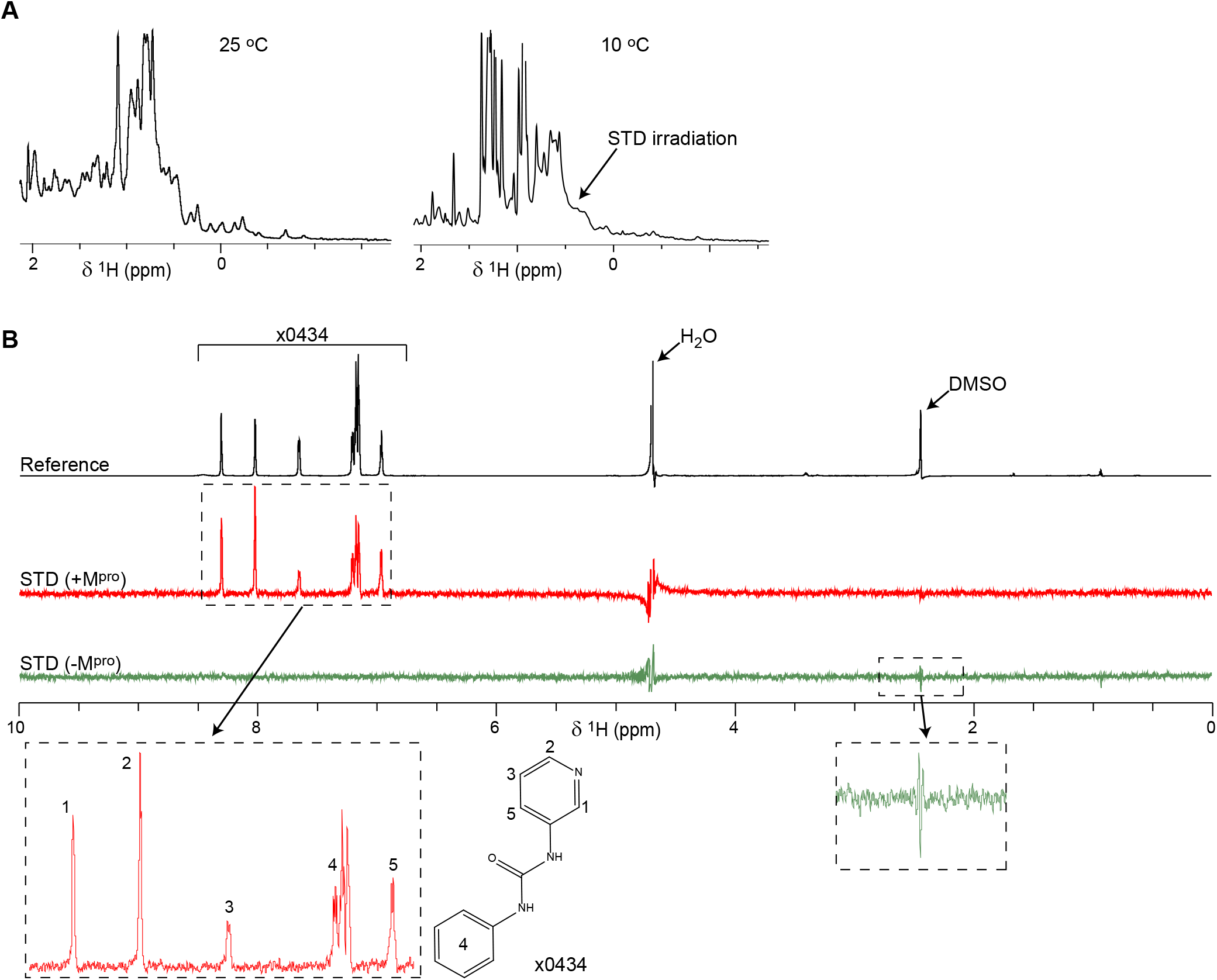
ID and STD-NMR spectra of SARS-CoV-2 M^pro^. A) Methyl regions from ^1^H NMR spectra of recombinant SARS-CoV-2 M^pro^. The spectrum on the left was recorded from a 10 μM protein concentration sample in a 5 mm NMR tube at 25 °C using an excitation sculpting water-suppression method (24). 512 acquisitions with recycle delay of 1.25 sec were averaged, for a total experiment time of just over 10 min. The spectrum on the right was recorded from a 10 μM M^pro^ sample in a 3 mm NMR tube at 10 °C, using the same pulse sequence and acquisition parameters. For both spectra, data were processed with a quadratic sine function prior to Fourier transformation. Protein resonances are weaker in the 10 °C spectrum due to lower temperature and the reduced amount of sample used for acquisition in the smaller NMR tube. The position where on-resonance irradiation was applied for STD spectra is indicated. B) Vertically offset ^1^H STD-NMR spectra from ligand x0434 binding to M^pro^. The reference spectrum is in black with the x0434, H_2_O and DMSO ^1^H resonances indicated. The STD spectrum of x0434 in the presence of M^pro^ is shown in red while that in the absence of M^pro^ is in green. STD spectra are scaled up 64x compared to the reference spectrum. Bottom panels correspond to magnified views of the indicated spectral regions, with x0434 resonances assigned to chemical groups of that ligand as shown.

### Ligand handling

Compounds for the initial STD-NMR assessment of crystallographic fragment binding to M^pro^ were provided by the XChem group at Diamond Light Source in the form of a 384-well plated library (DSI-poised, Enamine), with compounds dissolved in D_6_-DMSO at 500 mM nominal concentration. 1 μL of dissolved compounds was aspirated from this library and immediately mixed with 9 μL of D_6_-DMSO for a final fragment concentration of 50 mM, from which NMR samples were formulated. For titrations of the same crystallographic fragments compounds were procured directly from Enamine in the form of lyophilized powder, which was dissolved in D_6_-DMSO to derive compound stocks at 10 mM and 100 mM concentrations for NMR sample formulation.

STD-NMR assays of bespoke M^pro^ ligands used compounds commercially synthesised for COVID Moonshot. These ligands were provided to us by the XChem group in 96-well plates, containing 0.7 μL of 20 mM D_6_-DMSO-disolved compound per well. Plates were created using an Echo liquid handling robot (Labcyte) and immediately sealed and frozen at −20 °C. For use, ligand plates were thoroughly defrosted at room temperature and spun at 3,500 *g* for 5 minutes. In single-concentration STD-NMR experiments, 140 μL of a pre-formulated mixture of M^pro^ and NMR buffer with D_2_O and D_6_-DMSO were added to each well to create the final NMR sample. For STD-NMR competition experiments, 0.5 μL of ligands were aspirated from the plates and immediately mixed with 19.5 μL of D_6_-DMSO for final ligand concentration of 0.5 mM from which NMR samples were formulated.

### Molecular dynamics (MD) simulations

The monomeric complexes of M^pro^ bound to chemical fragments were obtained from the RCSB Protein Data Bank entries 5R81 (ligand x0195), 5REB (x0387), 5RGI (x0397), 5RGK (x0426), 5R83 (x0434) and 5REH (x0540) for MD simulations with GROMACS version 2018 (25) and the AMBER99SB-ILDN force field (26). All complexes were inserted in a pre-equilibrated box containing water implemented using the TIP3P water model (26). Force field parameters for the six ligands were generated using the general Amber force field and HF/6 – 31G* – derived RESP atomic charges (27). The reference system consisted of the protein, the ligand, ^~^31,400 water molecules, 95 Na and 95 Cl ions in a 100 x 100 x 100 Å simulation box, resulting in a total number of ^~^98,000 atoms. Each system was energy-minimized and subsequently subjected to a 20 ns MD equilibration, with an isothermal-isobaric ensemble using isotropic pressure control (28), and positional restraints on protein and ligand coordinates. The resulting equilibrated systems were replicated 4 times and independent 200 ns MD trajectories were produced with a time step of 2 fs, in constant temperature of 300 K, using separate v-rescale thermostats (28) for the protein, ligand and solvent molecules. Lennard-Jones interactions were computed using a cut-off of 10 Å and electrostatic interactions were treated using particle mesh Ewald (29) with the same real-space cut-off. Analysis on the resulting trajectories was performed using MDAnalysis (30, 31). Structures were visualised using PyMOL (32).

### Notes

The enzymatic inhibition potential of M^pro^ ligands, measured by RapidFire mass spectroscopy (17), was retrieved from the Collaborative Drug Discovery database (33).

## Results

### STD-NMR assays of M^pro^ ligand binding

M^pro^ forms dimers in crystals via an extensive interaction interface involving two domains (15). M^pro^ dimers likely have a sub-μM solution dissociation constant (K_d_) by analogy to previously studied 3C-like coronavirus proteases (34). At the 10 μM protein concentration of our NMR assays M^pro^ is, thus, expected to be dimeric with an estimated molecular weight of nearly 70 kDa. Despite the relatively large size of M^pro^ for solution NMR, ^1^H spectra of the protease readily showed the presence of multiple up-field shifted (<0.5 ppm) peaks corresponding to protein methyl groups (Fig. 1A). In addition to demonstrating that M^pro^ is folded under the conditions tested, these spectra allowed us to identify the chemical shifts of M^pro^ methyl groups that may be suitable for on-resonance irradiation in STD-NMR experiments. Trials with on-resonance irradiation applied to different methyl group peaks showed that irradiating at 0.5 ppm (Fig. 1A) produced the strongest STD signal from ligands in the presence of M^pro^, while simultaneously avoiding ligand excitation that would yield false-positive signals in the absence of M^pro^ (Fig. 1B). Further, we noted that small molecules abundant in the samples but not binding specifically to M^pro^, such as DMSO, produced pseudo-dispersive residual signal lineshapes in STD spectra, while true M^pro^ ligands produced peaks in STD with absorptive ^1^H lineshapes. We surmised that STD-NMR is suitable for screening ligand binding to M^pro^, requiring relatively small amounts (10-50 μgr) of protein and time (under 1 hour) per sample studied.

The strength of STD signal is quantified by calculating the ratio of integrated signal intensity of peaks in the STD spectrum over that of the reference spectrum (STD_ratio_). The STD_ratio_ factor is inversely proportional to ligand K_d_, as 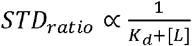 where [L] is ligand concentration. Measuring STD_ratio_ values over a range of ligand concentrations allows fitting of the proportionality constant and calculation of ligand K_d_. However, time and sample-amount considerations, including the limited availability of bespoke compounds synthesized for the COVID Moonshot project, made recording full STD-NMR titrations impractical for screening hundreds of ligands. Thus, we evaluated whether measuring the STD_ratio_ value at a single ligand concentration may be an informative alternative to K_d_, provided restraints could be placed, for example, on the proportionality constant.

Theoretical and practical considerations suggested that three parameters influence our evaluation of single-concentration STD_ratio_ values towards an affinity context. Firstly, the STD_ratio_ factor is affected by the efficiency of NOE magnetisation transfer between protein and ligand, which in turn depends on the proximity of ligand and protein groups, and the chemical nature of these groups (20–22). To minimize the influence of these factors across diverse ligands, we sought to quantify the STD_ratio_ of only aromatic ligand groups, and only consider those showing the strongest STD signal; thus, that are in closest proximity to the protein. Second, STD-NMR assays require ligand exchange between protein-bound and -free states in the timeframe of the experiment; strongly bound compounds that dissociate very slowly from the protein would yield reduced STD_ratio_ values compared to weaker ligands that dissociate more readily. Structures of M^pro^ with many different ligands show that the protein conformation does not change upon complex formation and that the active site is fully solvent-exposed (18), which suggests that ligand association can proceed with high rate (10^7^ – 10^8^ M^−1^s^−1^). Under this assumption, the ligand dissociation rate is the primary determinant of interaction strength. Given the duration of the STD-NMR experiment in our assays, and the ratios of ligand:protein used, we estimated that significant protein – ligand exchange will take place even for interactions as strong as low-μM K_d_. Finally, uncertainties or errors in nominal ligand concentration skew the correlation of STD_ratio_ to compound affinities; as shown in Fig. S1, STD_ratio_ values increase strongly when very small amounts of ligands are assessed. Thus, overly large STD_ratio_ values may be measured if ligand concentrations are significantly lower than anticipated.

### Quantitating M^pro^ binding of ligands identified by crystallographic screening

Mindful of the limitations inherent to measuring single-concentration STD_ratio_ values, and prior to using STD-NMR to evaluate bespoke M^pro^ ligands, we used this method to assess binding to the protease of small chemical fragments identified in crystallographic screening experiments (18). In crystallographic screening campaigns of other target proteins such fragments were seen to have very weak affinities (> 1 mM K_d_, e.g. (35)), thereby satisfying the exchange criterion set out above. 39 non-covalent M^pro^ interactors are part of the DSI-poised fragment library to which we were given access, comprising 17 active site binders, two compounds targeting the M^pro^ dimerisation interface and 20 molecules binding elsewhere on the protein surface (18). We initially recorded STD-NMR spectra from these compounds in the absence of M^pro^ to confirm that we obtained no or minimal STD signal when protease is omitted, and to verify ligand identity from reference ^1^H spectra. Five ligands gave no solution NMR signal or produced reference ^1^H spectra inconsistent with the compound chemical structure; these ligands were not evaluated further. Samples of 10 μM M^pro^ and 0.8 mM nominal ligand concentration were then formulated from the remaining 34 compounds (Table S1), and STD-NMR spectra were recorded, from which only aromatic ligand STD signals were considered for further analysis.

We observed large variations in STD signal intensity and STD_ratio_ values in the presence of M^pro^ across compounds (Fig. 2A,B; Table S1), with many ligands producing little or no STD signal, suggesting substantial differences in compound affinity for the protease. However, we also noted that ligand reference spectra different substantially in intensity (Fig. 2C), despite compounds being at the same nominal concentration. Integrating ligand peaks in these reference spectra revealed differences in per-^1^H intensity of up to ^~^15-fold, indicating significant variation of ligand concentrations in solution (Table S1). Such concentration differences could arise from errors in sample formulation or from concentration inconsistencies in the compound library. To evaluate the former we also integrated the residual ^1^H signal of D_6_-DMSO in our reference spectra, and found it to vary by less than 35% across any pair of samples (11% average deviation). As DMSO was added alongside ligands in our samples, we concluded that sample formulation may have contributed errors in compound concentration of up to ^~^1/3, but did not account for the ^~^15-fold differences in concentration observed.

**Figure 2:**
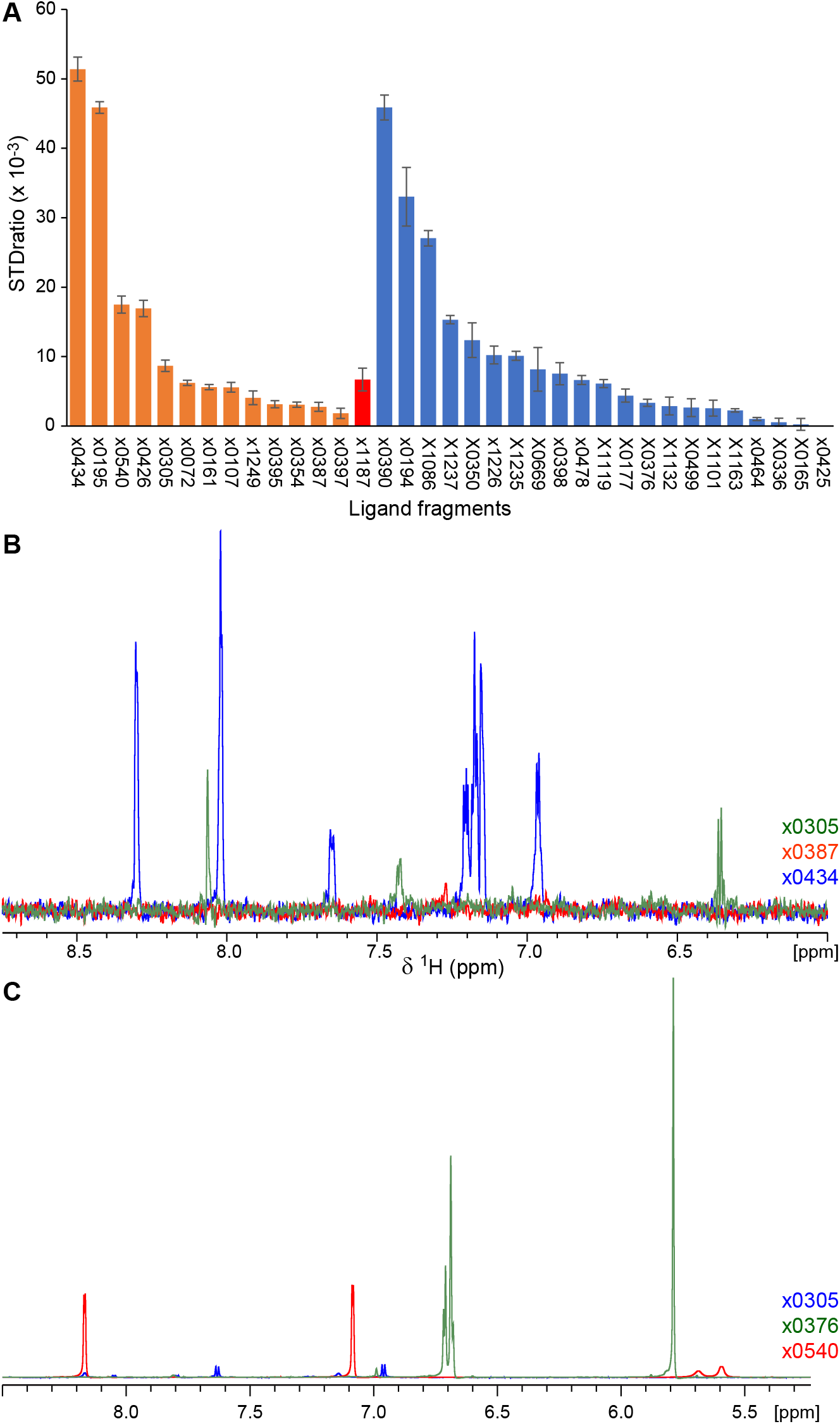
Assessment of fragment binding to M^pro^. A) STD_ratio_ values for chemical fragments identified by crystallographic screening as binding to M^pro^ (18). Ligands binding to the M^pro^ active site are coloured orange, at the M^pro^ dimer interface in red, and elsewhere on the protein surface in blue. B) Overlay of STD-NMR spectra from fragments x0305, x0387 and x434, which bind the M^pro^ active site, showing the ligand aromatic region in the presence of M^pro^. Spectra are colour coded per ligand as indicated. As seen, the three fragments yield significantly different STD signal intensities captured in the STD_ratio_ values shown in (A). C) Overlay of reference spectra from fragments x305, x376 and x540, showing the ligand aromatic region. Peak intensities vary substantially, suggesting significant differences in ligand concentration.

Given that differences in compound concentration can skew the relative STD_ratio_ values of ligands (Fig. S1), and that such concentration differences were also observed among newly designed M^pro^ inhibitors (see below), we questioned whether recording STD_ratio_ values under these conditions can provide useful information. To address this question we attempted to quantify the affinity of crystallographic fragments to M^pro^, selecting ligands that showed clear differences in STD_ratio_ values in the assays above and focusing on compounds binding at the M^pro^ active site; hence, that are of potential interest to inhibitor development. We performed M^pro^ binding titrations monitored by STD-NMR of compounds x0195, x0354, x0426 and x0434 in 50 μM – 4 mM concentrations (Fig. S2), and noted that only compounds x0434 and x0195, which show the highest STD_ratio_ (Fig. 2A), bound strongly enough for an affinity constant to be estimated (K_d_ of 1.6 ± 0.2 mM and 1.7 ± 0.2 mM, respectively). In contrast, the titrations of x0354 and x0426, which yielded lower STD_ratio_ values, could not be fit to extract a K_d_ indicating weaker binding to M^pro^.

To further this analysis, we assessed the binding of fragments x0195, x0387, x0397, x0426, x0434 and x0540 to the M^pro^ active site using quadruplicate atomistic molecular dynamics (MD) simulations of 200 nsec duration. As shown in Fig. S3A,B, and Movies S1 and S2, fragments with high STD_radio_ values (x0434 and x0195) always located in the M^pro^ active site despite exchanging between different binding conformations (Fig. S4), with average ligand root-mean-square-deviation (RMSD) of 3.2 Å and 5.1 Å respectively after the first 100 nsec of simulation. Medium STD_ratio_ value fragments (x0426 and x0540, Fig. S3C,D, and Movies S3 and S4) show average RMSDs of approximately 9 Å in the same simulation timeframe, frequently exchanging to alternative binding poses and with x0540 occasionally exiting the M^pro^ active site. In contrast, fragments showing very little STD NMR signal (x0397 and x0387, Fig. S3E,F, and Movies S5 and S6) regularly exit the M^pro^ active site and show average RMSDs in excess of 15 Å with very limited stability. Combining the quantitative K_d_ and MD information above, we surmised that, despite limitations inherent in this type of analysis and uncertainties in ligand amounts, STD_ratio_ values recorded at single compound concentration can act as proxy measurements of M^pro^ affinity for ligands.

### Assessment of M^pro^ binding by COVID Moonshot ligands

We proceeded to characterise by STD-NMR the M^pro^ binding of bespoke ligands created as part of the COVID Moonshot project and designed to act as non-covalent inhibitors of the protease (17). Similar to the assays of crystallographic fragments above, we focused our analysis of STD signals to aromatic moieties of ligands binding to the M^pro^ active side and extracted STD_ratio_ values only from the strongest STD peaks. Once again, we noted substantial differences in apparent compound concentrations, judging from reference ^1^H spectral intensities (Fig. 3A), which could not be attributed to errors in sample preparation as the standard deviation of residual ^1^H intensity in the D_6_-DMSO peak did not exceed 5% in any of the ligand batches tested. Crucially, out of 650 different molecules tested, samples of 35 compounds (7.6%) contained no ligand and 86 (13.2%) very little ligand (Fig. 3A). In these cases, NMR assays were repeated using a separate batch of compound; however, 96.2% of repeat experiments yielded the same outcome of no or very little ligand in the NMR samples.

**Figure 3:**
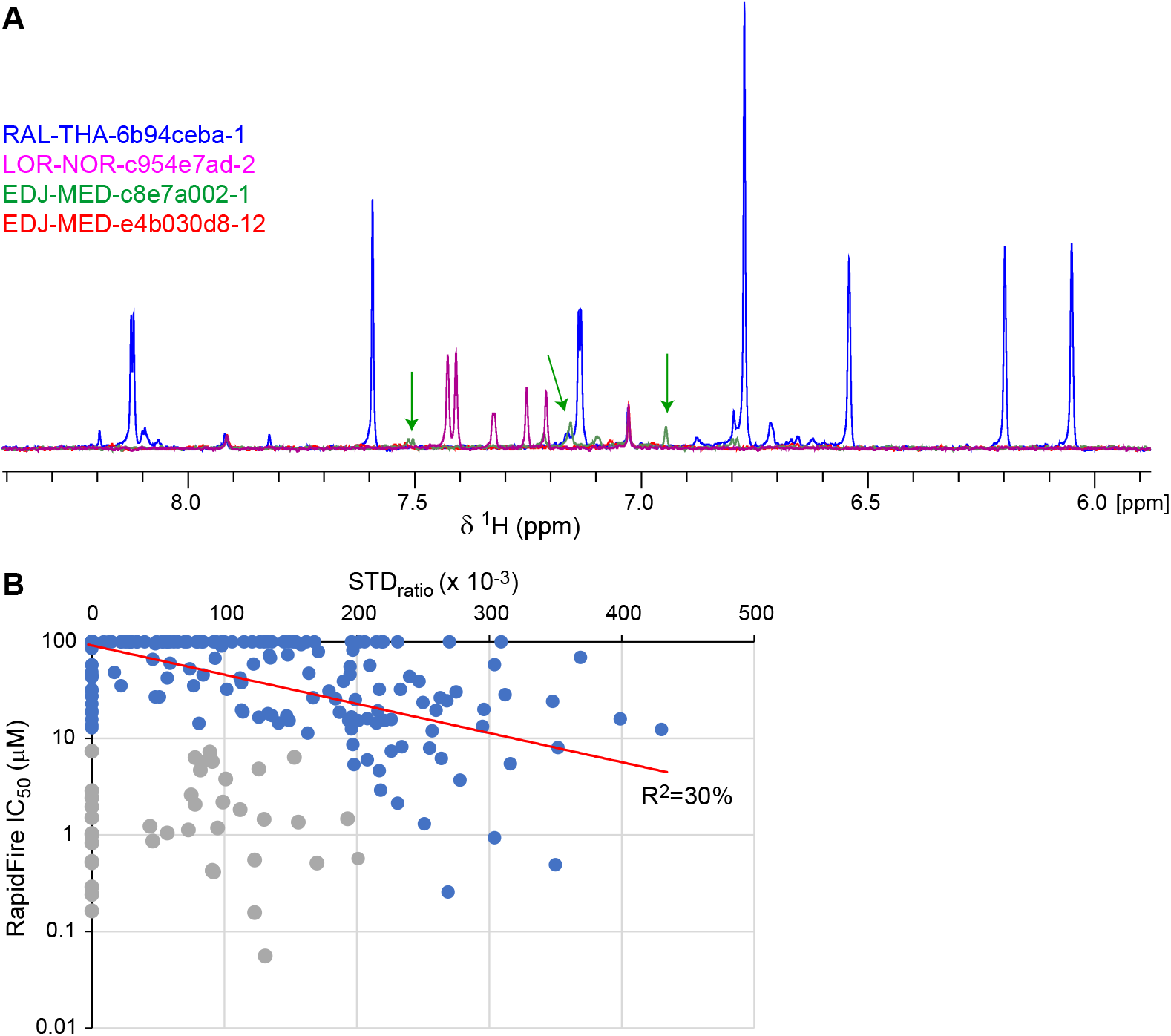
STD-NMR of COVID Moonshot ligands binding to M^pro^. A) Overlay of reference spectra from the indicated COVID Moonshot ligands, showing the ligand aromatic region in each case. in the presence of M^pro^. Spectra are colour coded per ligand as indicated. As seen, peak intensities vary substantially, suggesting significant differences in ligand concentration. Peaks of ligand EDJ-MED-c8e7a002-1 (green) are indicated by arrows; ligand EDJ-MED-e4b030d8-12 (red) produced no peaks in the NMR spectrum. B) Plot of STD_ratio_ values from COVID Moonshot ligands assessed by STD-NMR against their IC_50_ value estimated by RapidFire mass spectroscopy enzymatic assays (17). Ligands in blue show weak correlation between the two methods (red line, corresponding to an exponential function along the IC50 dimension). Ligands in grey represent outliers of the STD-NMR or enzymatic method as discussed in the text.

We measured STD_ratio_ values from samples were ligands produced sufficiently strong reference ^1^H NMR spectra to be readily visible, and deposited these values and associated raw NMR data to the Collaborative Drug Discovery database (33). Some of these ligands were assessed independently for enzymatic inhibition of M^pro^ using a mass spectroscopy method as part of the COVID Moonshot collaboration (17). Where both parameters are available, we compared the STD_ratio_ values and 50% inhibition concentrations (IC_50_) of these ligands. As shown in Fig. 3B, STD_ratio_ and IC_50_ values show weak correlation (R^2^=30%) for most ligands tested; however, a subset of ligands displayed conspicuously low or even no STD signals considering their effect on M^pro^ activity, and presented themselves as outliers in the correlation graph. As these outlier ligands had IC_50_ values below 10 μM, suggesting that their affinities to the protease may be in the μM K_d_ region, we considered whether our approach gives rise to false-negative STD results, for example through slow ligand dissociation from M^pro^.

To address this question, we derived an assay whereby the bespoke, high-affinity M^pro^ inhibitor would outcompete a lower-affinity ligand known to provide strong STD signal from the protease active site. In these experiments the lower-affinity ligand would act as ‘spy’ molecule whose STD signal reduces as function of inhibitor concentration. We used fragment x0434, which yields substantial STD signal with M^pro^ (Fig. 1B and 2A), as ‘spy’, and tested protease inhibitors EDJ-MED-a364e151-1, LON-WEI-ff7b210a-5, CHO-MSK-6e55470f-14 and LOR-NOR-30067bb9-11 as x0434 competitors. Of these inhibitors, EDJ-MED-a364e151-1 gave rise to substantial STD signal in earlier assays, whereas the remaining produced little or no STD signal; yet, all four inhibitors were reported to have low-μM or sub-μM IC50 values based on M^pro^ enzymatic assays. In these competition experiments, both EDJ-MED-a364e151-1 and LON-WEI-ff7b210a-5 yielded K_d_ parameters comparable to the reported IC_50_ values (Fig. S5A,B), showing that at least in the case of LON-WEI-ff7b210a-5 the absence of STD signal in the single-concentration NMR assays above represented a false-negative result. In contrast, CHO-MSK-6e55470f-14 and LOR-NOR-30067bb9-ll were unable to compete x0434 from the protease active site (Fig. S5C,D), suggesting that in these two cases the reported IC_50_ values do not reflect inhibitor binding to the protease, and that the weak STD signal of the initial assays was a better proxy of affinity. We surmised that although some low STD_ratio_ values of M^pro^ inhibitors may not accurately reflect compound affinity to the protease, such values cannot be discounted as a whole as they may correspond to non-binding ligands.

## Discussion

Fragment-based screening is a tried and tested method for reducing the number of compounds that need to be assessed for binding against a specific target in order to sample chemical space (36). Combined with X-ray crystallography, which provides information on the target site and binding pose of ligands, initial fragments can quickly be iterated into potent and specifically-interacting compounds. The COVID Moonshot collaboration (17) took advantage of crystallographic fragment-based screening (18) to initiate the design of novel inhibitors targeting the essential main protease of the SARS-CoV-2 coronavirus; however crystallographic structures do not report on ligand affinity and inhibitory potency in enzymatic assays does not always correlate with ligand binding. Thus, supplementing these methods with solution NMR tools highly sensitive to ligand binding can provide a powerful combination of orthogonal information and assurance against false starts.

We showed that STD-NMR is a suitable method for characterising ligand binding to M^pro^, allowing us to assess ligand interactions using relatively small amounts of protein and in under one hour of experiment time per ligand (Fig. 1B). However, screening compounds in a high-throughput manner is not compatible with the time- and ligand-amount requirements of full STD-NMR titrations. Thus, we resorted to using an unconventional metric, the single-concentration STD_ratio_ value, as proxy for ligand affinity. Although this metric has limitations due to its dependency on magnetisation transfer between protein and ligand, and on relatively rapid exchange between the ligand-free and -bound states, we demonstrated that it can nevertheless be informative. Specifically, the relative STD_ratio_ values of chemical fragments bound to the M^pro^ active site provided insight on fragment affinity (Fig. 2A), as crosschecked by quantitative titrations (Fig. S2) and MD simulations (Fig. S3). Furthermore, STD_ratio_ values of COVID Moonshot compounds held a weak correlation to enzymatic IC_50_ parameters (Fig. 3B), although false-negative and -positive results from both methods contribute to multiple outliers. Thus, in our view the biggest limitation of using the single-concentration STD_ratio_ value as metric relates to its supra-linear sensitivity to ligand concentration (Fig. S1), which as demonstrated here can vary substantially across ligands in a large project (Fig. 3A).

How then should the STD data recorded as part of COVID Moonshot be used? Firstly, we showed that at least for some bespoke M^pro^ ligands the STD_ratio_ value obtained is a better proxy for compound affinity compared to IC_50_ parameters from enzymatic assays (Fig. S5). This, inherently, is the value of employing orthogonal methods thereby minimizing the number of potential false results. Thus, when one is considering existing M^pro^ ligands to base the design of future inhibitors, a high STD_ratio_ value as well as low IC_50_ parameters are both desirable. Second, due to the aforementioned limitations of single-concentration STD_ratio_ value as proxy of affinity, and the influence of uncertainties in ligand concentrations, we believe that comparisons of compounds and derivatives differing by less than ^~^50% in STD_ratio_ is not meaningful. Rather, we propose that the STD_ratio_ values of M^pro^ ligands measured and available at the CDD database should be treated as a qualitative metrics of compound affinity.

In conclusion, we presented here protocols for the assessment of SARS-CoV-2 M^pro^ ligands using STD-NMR spectroscopy, and evaluated the relative qualitative affinities of chemical fragments and compounds designed as part of COVID Moonshot. Although development of novel antivirals to combat COVID-19 is still at an early stage, we hope that this information will prove valuable to groups working towards such treatments.

## Acknowledgements

We are grateful to Nick Soffe for maintenance of the Oxford Biochemistry solution NMR facility, to Claire Strain-Damerell, Petra Lukacik and Martin A. Walsh for advice on M^pro^ production, to Anthony Aimon and Frank von Delft for providing the DSI-poised fragment library, to Adrián García, Nil Casajuana and Clàudia Llinàs del Torrent for advice with MD analysis tools, and to Leonardo Pardo for providing access to high-performance computing facilities. This work was supported by philanthropic donations to the University of Oxford COVID-19 Research Response Fund and the Oxford Glycobiology Institute Endowment. The Oxford Biochemistry NMR facility was supported by the Wellcome Trust (094872/Z/10/Z), the Engineering and Physical Sciences Research Council (EP/R029849/1), the Wellcome Institutional Strategic Support Fund, the EPA Cephalosporin Fund and the John Fell OUP Research Fund. This work was also supported by the “Reinforcement of Postdoctoral Researchers - 2nd Cycle” (MIS-5033021), implemented by the Greek State Scholarships Foundation (IKY).

## References

1. WHO. Coronavirus disease 2019 [Available from: https://www.who.int/emergencies/diseases/novel-coronavirus-2019.

2. Kucharski AJ, Russell TW, Diamond C, Liu Y, Edmunds J, Funk S, et al. Early dynamics of transmission and control of COVID-19: a mathematical modelling study. Lancet Infect Dis. 2020;20(5):553–8.

3. Wu F, Zhao S, Yu B, Chen YM, Wang W, Song ZG, et al. A new coronavirus associated with human respiratory disease in China. Nature. 2020;579(7798):265–9.

4. Zhu N, Zhang D, Wang W, Li X, Yang B, Song J, et al. A Novel Coronavirus from Patients with Pneumonia in China, 2019. N Engl J Med. 2020;382(8):727–33.

5. Bermingham A, Chand MA, Brown CS, Aarons E, Tong C, Langrish C, et al. Severe respiratory illness caused by a novel coronavirus, in a patient transferred to the United Kingdom from the Middle East, September 2012. Euro Surveill. 2012;17(40):20290.

6. Kuiken T, Fouchier RA, Schutten M, Rimmelzwaan GF, van Amerongen G, van Riel D, et al. Newly discovered coronavirus as the primary cause of severe acute respiratory syndrome. Lancet. 2003;362(9380):263–70.

7. Zaki AM, van Boheemen S, Bestebroer TM, Osterhaus AD, Fouchier RA. Isolation of a novel coronavirus from a man with pneumonia in Saudi Arabia. N Engl J Med. 2012;367(19):1814–20.

8. Thiel V, Ivanov KA, Putics A, Hertzig T, Schelle B, Bayer S, et al. Mechanisms and enzymes involved in SARS coronavirus genome expression. J Gen Virol. 2003;84(Pt 9):2305–15.

9. Bredenbeek PJ, Pachuk CJ, Noten AF, Charite J, Luytjes W, Weiss SR, et al. The primary structure and expression of the second open reading frame of the polymerase gene of the coronavirus MHV-A59; a highly conserved polymerase is expressed by an efficient ribosomal frameshifting mechanism. Nucleic Acids Res. 1990;18(7):1825–32.

10. Hilgenfeld R. From SARS to MERS: crystallographic studies on coronaviral proteases enable antiviral drug design. FEBS J. 2014;281(18):4085–96.

11. Ghosh AK, Xi K, Grum-Tokars V, Xu X, Ratia K, Fu W, et al. Structure-based design, synthesis, and biological evaluation of peptidomimetic SARS-CoV 3CLpro inhibitors. Bioorg Med Chem Lett. 2007;17(21):5876–80.

12. Verschueren KH, Pumpor K, Anemuller S, Chen S, Mesters JR, Hilgenfeld R. A structural view of the inactivation of the SARS coronavirus main proteinase by benzotriazole esters. Chem Biol. 2008;15(6):597–606.

13. Yang H, Yang M, Ding Y, Liu Y, Lou Z, Zhou Z, et al. The crystal structures of severe acute respiratory syndrome virus main protease and its complex with an inhibitor. Proc Natl Acad Sci U S A. 2003;100(23):13190–5.

14. Yang H, Xie W, Xue X, Yang K, Ma J, Liang W, et al. Design of wide-spectrum inhibitors targeting coronavirus main proteases. PLoS Biol. 2005;3(10):e324.

15. Zhang L, Lin D, Sun X, Curth U, Drosten C, Sauerhering L, et al. Crystal structure of SARS-CoV-2 main protease provides a basis for design of improved alpha-ketoamide inhibitors. Science. 2020;368(6489):409–12.

16. Rut W, Groborz K, Zhang L, Sun X, Zmudzinski M, Pawlik B, et al. SARS-CoV-2 M(pro) inhibitors and activity-based probes for patient-sample imaging. Nat Chem Biol. 2020.

17. Achdout H, Aimon A, Bar-David E, Barr H, Ben-Shmuel A, et al. COVID Moonshot: Open Science Discovery of SARS-CoV-2 Main Protease Inhibitors by Combining Crowdsourcing, High-Throughput Experiments, Computational Simulations, and Machine Learning. bioRxiv. 2020.

18. Douangamath A, Fearon D, Gehrtz P, Krojer T, Lukacik P, Owen CD, et al. Crystallographic and electrophilic fragment screening of the SARS-CoV-2 main protease. Nat Commun. 2020;11(1):5047.

19. El-Baba TJ, Lutomski CA, Kantsadi AL, Malla TR, John T, Mikhailov V, et al. Allosteric Inhibition of the SARS-CoV-2 Main Protease: Insights from Mass Spectrometry Based Assays. Angew Chem Int Edit. 2020.

20. Mayer M, Meyer B. Characterization of Ligand Binding by Saturation Transfer Difference NMR Spectroscopy. Angew Chem Int Ed Engl. 1999;38(12):1784–8.

21. Becker W, Bhattiprolu KC, Gubensak N, Zangger K. Investigating Protein-Ligand Interactions by Solution Nuclear Magnetic Resonance Spectroscopy. Chemphyschem. 2018;19(8):895–906.

22. Walpole S, Monaco S, Nepravishta R, Angulo J. STD NMR as a technique for ligand screening and structural studies. Methods in Enzymology. 615: Elsevier; 2019. p. 423–51.

23. Rogala KB, Dynes NJ, Hatzopoulos GN, Yan J, Pong SK, Robinson CV, et al. The Caenorhabditis elegans protein SAS-5 forms large oligomeric assemblies critical for centriole formation. Elife. 2015;4:e07410.

24. Hwang TL, Shaka AJ. Water Suppression That Works -Excitation Sculpting Using Arbitrary Wave-Forms and Pulsed-Field Gradients. Journal of Magnetic Resonance Series A. 1995;112(2):275–9.

25. Abraham MJ, Murtola T, Schulz R, Páll S, Smith JC, Hess B, et al. GROMACS: High performance molecular simulations through multi-level parallelism from laptops to supercomputers. SoftwareX. 2015;1:19–25.

26. Lindorff-Larsen K, Piana S, Palmo K, Maragakis P, Klepeis JL, Dror RO, et al. Improved side-chain torsion potentials for the Amber ff99SB protein force field. Proteins. 2010;78(8):1950–8.

27. Bayly CI, Cieplak P, Cornell W, Kollman PA. A well-behaved electrostatic potential based method using charge restraints for deriving atomic charges: the RESP model. J Phys Chem. 1993;97(40): 10269–80.

28. Bussi G, Zykova-Timan T, Parrinello M. Isothermal-isobaric molecular dynamics using stochastic velocity rescaling. J Chem Phys. 2009;130(7):074101.

29. Darden T, York D, Pedersen L. Particle mesh Ewald: An N· log (N) method for Ewald sums in large systems. J Chem Phys. 1993;98(12):10089–92.

30. Michaud-Agrawal N, Denning EJ, Woolf TB, Beckstein O. MDAnalysis: a toolkit for the analysis of molecular dynamics simulations. J Comput Chem. 2011;32(10):2319–27.

31. Gowers RJ, Linke M, Barnoud J, Reddy TJE, Melo MN, Seyler SL, et al., editors. MDAnalysis: A Python package for the rapid analysis of molecular dynamics simulations. 15th Python in Science Conference; 2016; Austin, TX.

32. DeLano WL. The PyMOL Molecular Graphics System. DeLano Scientific, San Carlos, CA, USA. http://www.pymol.org.2002.

33. Collaborative Drug Discovery database public access 2020 [Available from: https://www.collaborativedrug.com/public-access/.

34. Grum-Tokars V, Ratia K, Begaye A, Baker SC, Mesecar AD. Evaluating the 3C-like protease activity of SARS-Coronavirus: recommendations for standardized assays for drug discovery. Virus Res. 2008;133(1):63–73.

35. Davies TG, Wixted WE, Coyle JE, Griffiths-Jones C, Hearn K, McMenamin R, et al. Monoacidic Inhibitors of the Kelch-like ECH-Associated Protein 1: Nuclear Factor Erythroid 2-Related Factor 2 (KEAP1:NRF2) Protein-Protein Interaction with High Cell Potency Identified by Fragment-Based Discovery. J Med Chem. 2016;59(8):3991–4006.

36. Erlanson DA, Fesik SW, Hubbard RE, Jahnke W, Jhoti H. Twenty years on: the impact of fragments on drug discovery. Nat Rev Drug Discov. 2016;15(9):605–19.

