## Supplemental figures and captions for "A COVID Moonshot: assessment of ligand binding to the SARS-CoV-2 main protease by saturation transfer difference NMR spectroscopy"

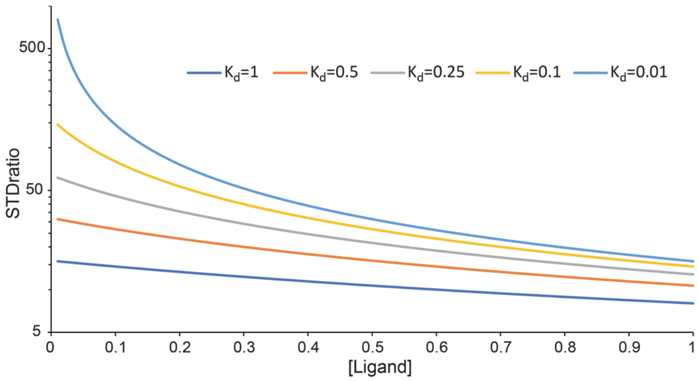


*Supplemental Figure 1: $STD_{ratio}$ values vary as function of ligand $K_d$ and concentration.* Shown here are simulations of $STD_{ratio}$ values for arbitrary ligand concentration (0 to 1) as function of the ligand interaction strength ($K_d$) expressed in the same concentration scale. As seen, strongly interacting ligands ($K_d$ of 0.1 to 0.01) produce overly large $STD_{ratio}$ values if in low concentration, thereby skewing the correlation of $STD_{ratio}$ values to ligand affinity.

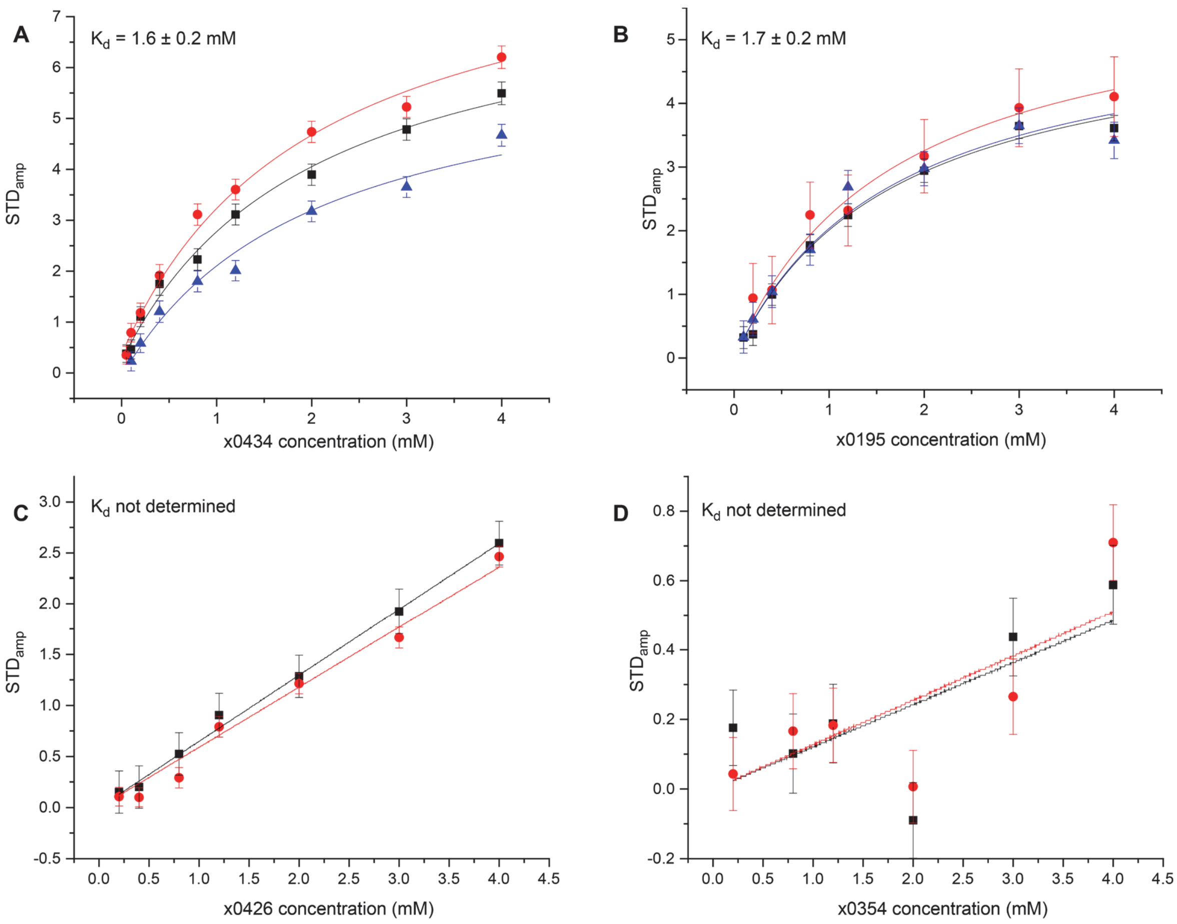


*Supplemental Figure 2: Quantification of interaction affinities for chemical fragments binding to the $M^{pro}$ active site.* Shown here are plots of STD amplification factors ($STD_{amp}$, equal to $STD_{ratio}$ multiplied by ligand excess) versus fragment concentration, measured in STD-NMR experiments of fragments with $M^{pro}$. Each data series within panels A-D corresponds to $STD_{amp}$ values derived from separate resonance peaks of the indicated ligands. Data were fit to a single-site association model with global $K_d$ for each ligand. As seen, whereas ligands x0434 and x0195 produced plots that could be fit to extract $K_d$ values, the affinities of ligands x0426 and x0354 could not be determined.

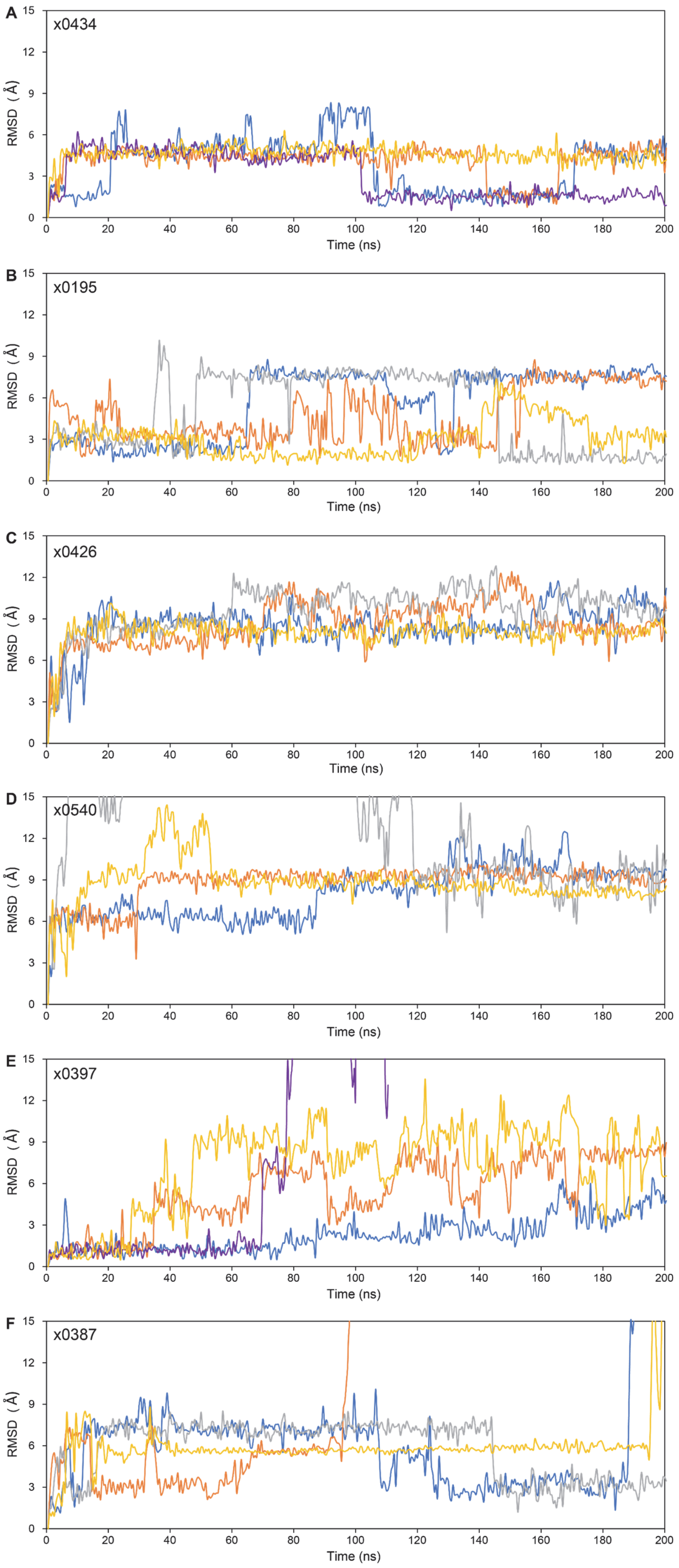
A
x0434
B
x0195
C
x0426
D
x0540
E
x0397
F
x0387
RMSD (Å)
0
3
6
9
12
15
Time (ns)
0
20
40
60
80
100
120
140
160
180
200

*Supplemental Figure 3: MD simulations of chemical fragments binding to the $M^{pro}$ active site.* Shown here are plots of ligand RMSD from the simulation starting point as function of simulation time. Each simulation was performed four times as indicated by the different colour traces.

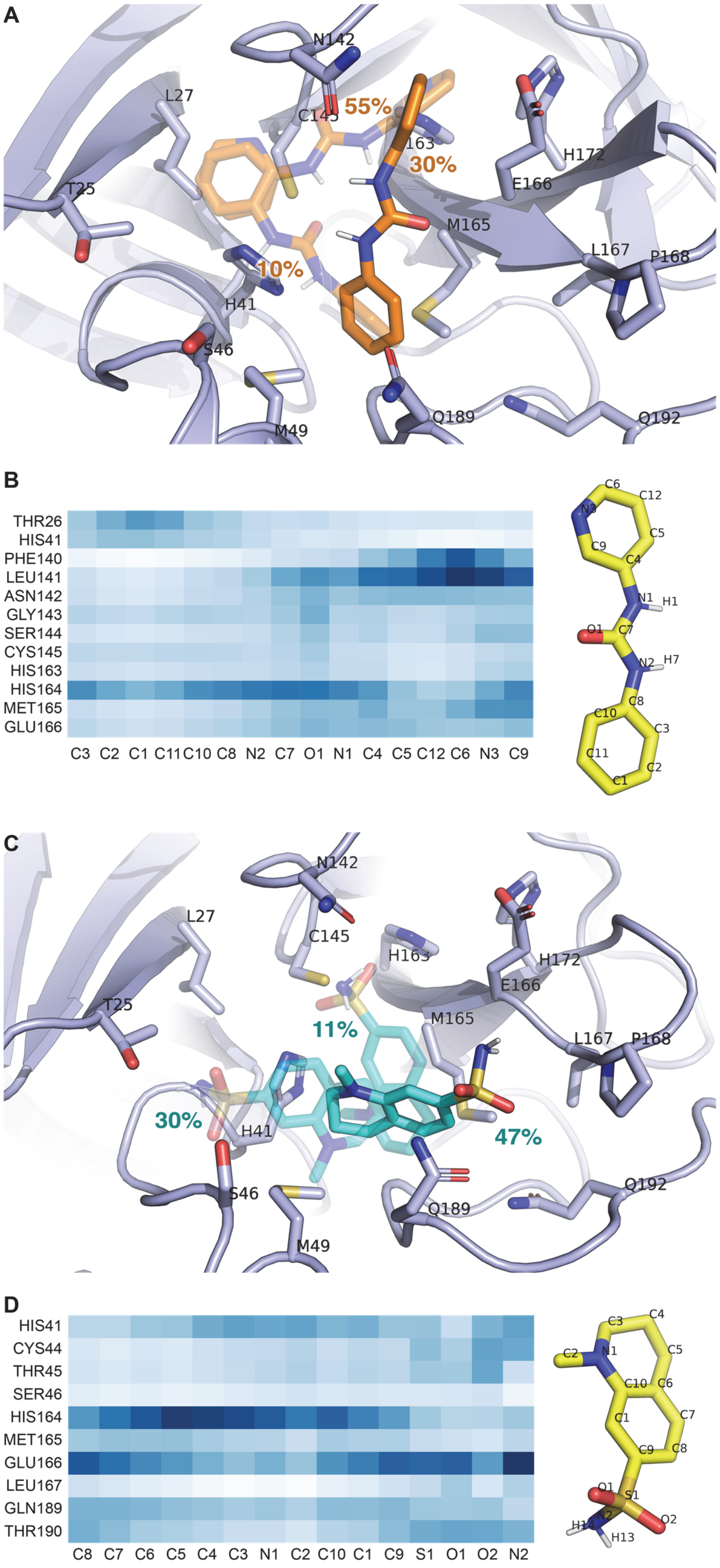


*Supplemental Figure 4: Binding poses of compounds x0434 and x0195 in the M$^{pro}$ active site.* A) The M$^{pro}$ active site with ligand x0434 (orange sticks) shown in the three most prevalent binding poses adopted during the quadruplicate MD simulations. The relative populations of each pose are indicated. B) Contact probability map of M$^{pro}$ amino acids and x0434 atoms in the MD simulations. The x0434 structure and atom assignments are shown on the right. Contact probabilities have been normalized to the most prevalent pair (darker blue shades indicating higher probability). C,D) Similar representations of the $M^{pro}$ active site in simulations of ligand x0195 (C) and contact probability map of this ligand (D).

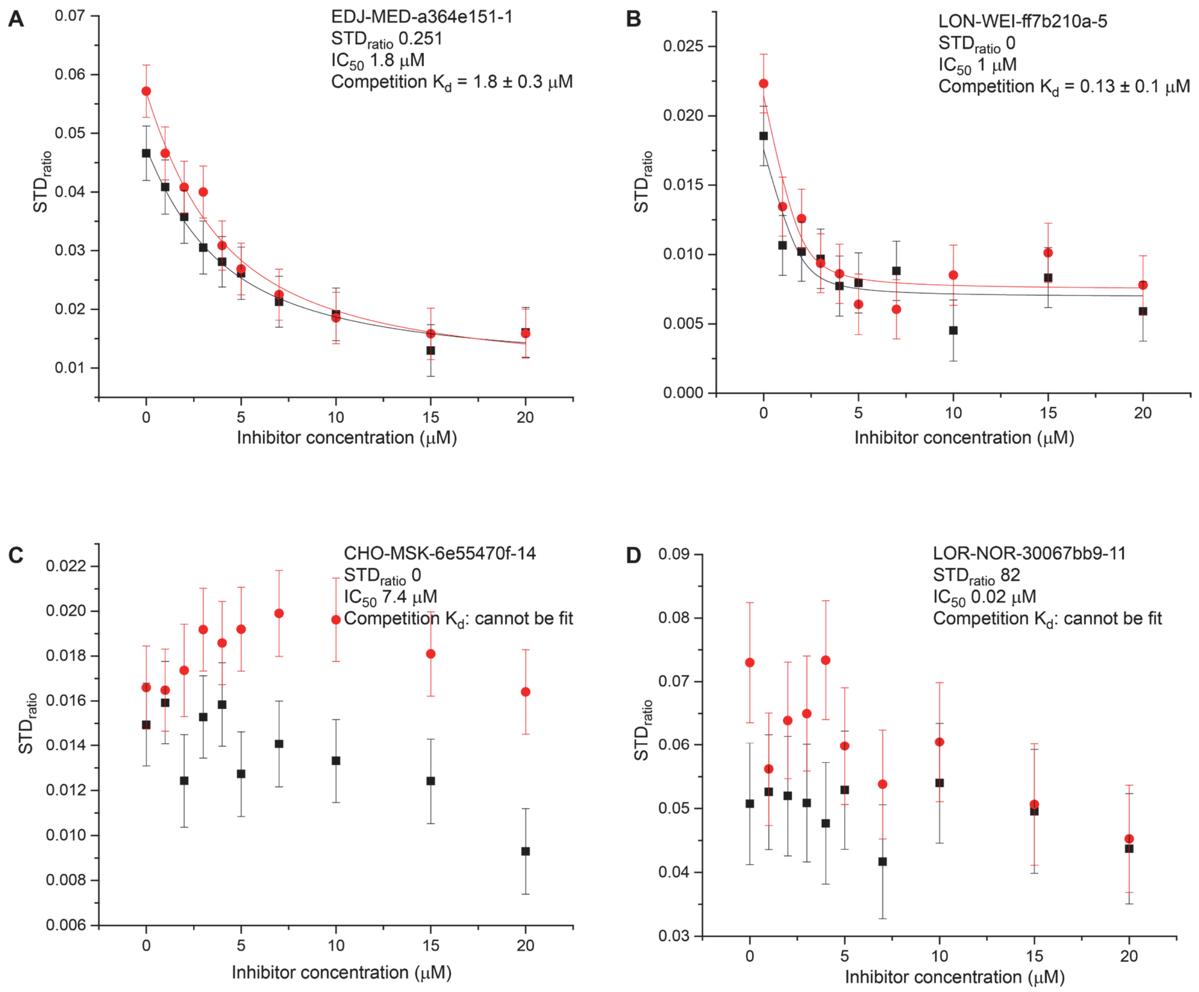


*Supplemental Figure 5: Quantification of interaction affinities for COVID Moonshot ligands binding to $M^{pro}$.* Shown here are plots of $STD_{ratio}$ values derived from two NMR peaks of fragment x0434 as function of a competing COVID Moonshot ligand concentration. For each ligand, the $STD_{ratio}$ values derived from single-point assays and the $IC_{50}$ values from RapidFire enzymatic assays are indicated. The titrations of COVID Moonshot ligands EDJ-MED-a364e151-1 and LON-WEI-ff7b210a-5 could be fit to an ideal competition model to derive binding affinities for these compounds; in contrast, ligands CHO-MSK-6e55470f-14 and LOR-NOR-30067bb9-11 failed to displace fragment x0434 from the $M^{pro}$ active site in a significant manner.

*Supplemental Movies S1-S6. MD simulations of chemical fragments binding to the $M^{pro}$ active site.* Movie S1 is of fragment x0434; movie S2 of fragment x0195; movie S3 of fragment x0426; movie S4 of fragment x0540; movie S5 of fragment x0397; movie S6 of fragment x0387). $M^{pro}$ is shown in schematic representation and ligands as sticks. Hydrogen bonds are indicated by dashed yellow lines. Each pane corresponds to an independent simulation.
